# Development of three-dimensional articular cartilage construct using silica nano-patterned substrate

**DOI:** 10.1101/472332

**Authors:** In-Su Park, Ye Ji Choi, Bo Ram Song, Hyo-Sop Kim, Sang-Hyug Park, Byung Hyune Choi, Jae-Ho Kim, Byoung-Hyun Min

**Affiliations:** Cell Therapy Center, Ajou University Medical Center, Suwon, Korea; Department of Physiology, Inha University College of Medicine, Incheon, Korea; Department of Molecular Science & Technology, Ajou University, Suwon, Korea; Department of Biomedical Engineering, Pukyong National University, Korea; Department of Orthopedic Surgery, Ajou University School of Medicine, Suwon, Korea

**Keywords:** Microenvironment, Three-dimensional cell aggregation, Silica nano-patterned substrates, Fetal cartilage-derived progenitor cells

## Abstract

Current strategies for cartilage cell therapy are mostly based on the use of autologous chondrocytes or mesenchymal stem cells (MSCs). However, these cells have limitations of a small number of cells available and of low chondrogenic ability, respectively. Many studies now suggest that fetal stem cells are more plastic than adult stem cells and can therefore more efficiently differentiate into target tissues. This study introduces, efficiency chondrogenic differentiation of fetal cartilage-derived progenitor cells (FCPCs) to adult cells can be achieved using a three-dimensional (3D) spheroid culture method based on silica nanopatterning techniques. In evaluating the issue of silica nano-particle size (Diameter of 300, 750, 1200 nm), each particle size was coated into the well of a 6-well tissue culture plate. FCPCs (2 x 10^5^ cells/well in 6-well plate) were seeded in each well with chondrogenic medium. In this studys, the 300 nm substrate that formed multi-spheroids and the 1200 nm substrate that showed spreading were due to the cell-cell adhesion force(via N-cadherin) and cell-substrate(via Integrin) force, the 750 nm substrate that formed the mass-aggregation can be interpreted as the result of cell monolayer formation through cell-substrate force followed by cell-cell contact force contraction. We conclude that our 3D spheroid culture system contributes to an optimization for efficient differentiation of FCPC, offers insight into the mechanism of efficient differentiation of engineered 3D culture system, and has promise for wide applications in regeneration medicine and drug discovery fields.

## Introduction

The self-repair of articular cartilage (AC) is difficult when it becomes injured due to its low regenerative capacity caused by the lack of blood supply, low cellularity, and a limited number of progenitor cells [1]. Over the last years, many studies have utilized mesenchymal stem cells (MSCs) isolated from various tissues, such as bone marrow, synovial membrane, fat, or cord blood [1]. However, MSCs have also been shown to have limited capacity to differentiate fully into chondrocytes and form intact cartilage tissues both in vitro and in vivo [2]. Meanwhile, a number of studies have shown that stem cells or progenitor cell derived from human fetal tissue is an encouraging cell source for cell therapy and tissue engineering efforts [3]. In our previous study, we proved the potential of FCPCs as a novel cell source for cartilage engineering. These FCPCs showed high cell yields, proliferating, multi-lineage differentiation ability [4]. These results suggest that FCPCs can serve as a suitable cell source for tissue engineering. Previously, several studies attempted to use fetal cartilage-derived cells in cartilage tissue engineering.

However, the two-dimensional (2D) culture method has important limitations in controlling stem cell differentiation pathways resulting in low differentiation efficiency [5]. To overcome these limitations of the 2D culture, three-dimensional (3D) culture is used as culture condition for cell microenvironment [6],[7]. Fuchs et al. reported that ovine fetal cartilage cells formed better cartilage tissue than adult chondrocytes by producing more matrix molecules in the pellet culture. This 3D culture used to enhanced differentiation marker gene [8],[9] and anti-inflammatory cytokine expression [10],[11] and stimulation of cellular ECM secretion [11],[12]. Cell-cell and cell-extracellular matrix (ECM) interactions are crucial for maintaining cell phenotype and for inducing effective differentiation in stem cells. 3D cell culture methods facilitate greater cell-cell and cell-ECM interactions, allowing cells to create an “in vivo-like” microenvironment and better preserve stem cell phenotype [13] and induce re-differentiation [14].

The cell behavior, such as morphology and adhesion, differentiation, is influenced by their surrounding microenvironment [15]. This cellular microenvironment is controlled by surface property cues through topographical, mechanical, chemical control [16]. Especially, A number of studies have topographic patterns for influencing cellular behavior, various Nano-post density triggered changes in cell cytoskeletal structure and differentiation [17], patterns made from Nanosize particles of various diameter was restricted cell spreading [18], a Nano-periodic patterns were significantly increasing the mechanical property in the self-assembled tissue through cytoskeleton alignment [19]. Based on a these studies, the recent paradigm of attempt induces spheroids formation as 3D culture through patterns substrate was reported [20]. The aggregation formation process, meanwhile, involves changes in cell morphology, indicative of the reorganization of the cytoskeleton. The dynamics of the actin cytoskeleton are induced through integrin-mediated cell-material interaction and cadherin-mediated cell-cell interactions [21],[22]. Simply, controlling the cell adhesion molecule enables to regulate the formation cell 3D aggregation.

This study used to fetal cartilage-derived progenitor cells (FCPCs), an important cell in the aggregation process and Nano-patterned substrate that able to control cell adhesion, for manufacture of three-dimensional cartilage constructs. Silica beads are well known by their stability in chemical structure and shape. We investigated for the effects of FCPCs aggregation formation on Nano-patterned substrate through particles that 300-1200nm in diameter. By examining FCPCs morphology and relative adhesion force, adhesion molecule, also, total volume and chondrogenic potential of various 3D constructs. We herein test hypothesis that change of FCPCs adhesive by using Nano-patterned substrate through surface particle-size will be induced 3D aggregation formation. Based on them, we manufactured a three-dimensional cartilage construct and analyzed this phenomenon. These 3D cartilage-like tissues using a 3D aggregation culture system, will be useful for in vitro studies of cartilage tissue engineering and cell therapy [23].

## Materials and Methods

### Silica bead synthesis

Monodispersed silica beads were synthesized using the Stober method as described in our previous report. The synthesized silica beads with an average diameter of 750 nm were purified by centrifugation and redispersed in ethanol [24].

### Cell Isolation and Culture

The study was approved by the institutional review board (IRB) of the Ajou University Medical Center (AJIRB-CRO-07-139) and was carried out with the written consent of all donors. Human fetal cartilage tissues (*n* = 2, F12w-c, M11w) were obtained from patients following elective termination at 12 weeks after gestation, and cells were isolated from the femoral head of the cartilage tissue. Cartilage tissues were cut into small pieces and treated with 0.1% collagenase type Π (Worthington Biochemical Corp, Freehold, NJ, USA) in high-glucose Dulbecco’s modified Eagle medium (DMEM; Hyclone, Logan, UT, USA) containing 1% fetal bovine serum(FBS; Biotechnics research, Inc.) at 37°C under 5% CO2. After 12h, isolated cells were cultured in DMEM supplemented with 10% FBS, Penicillin streptomycin (100 U/ml penicillin G (Gibco BRL), 100 μg/ml streptomyocin (Gibco BRL)), 5 ng/ml basic FGF (R&D systems, Recombinant human FGF basic146aa, USA). Cells were passaged at 80% confluence, where the plating density was approximately 8 × 10^3^ cells/cm^2^.

### Cell culture on Nano patterned substrate

In evaluating the issue of Nano-particle size (Diameter of 300, 750, 1200 nm), each particle size was coated into the well of a 6-well tissue culture plate. FCPCs (2 x 10^5^ cells/well in 6-well plate) were seeded in each well with chondrogenic medium. The chondrogenic medium consisted of DMEM supplemented with Penicillin streptomycin (100 U/ml penicillin G (Gibco BRL), 100 μg/ml streptomycin (Gibco BRL)), ITS supplement (Gibco BRL, 1.0 mg/ml insulin from bovine pancreas, 0.55 mg/ml human transferrin, and 0.5 mg/ml sodium selenite), 50 μg/ml ascorbic acid, 100 nM dexamethasone, 40 μg/ml proline, 1.25mg/ml bovine serum albumin (BSA), and 100 μg/ml sodium pyruvate (all from Sigma, ST. Louis, MO, USA).

The morphology of cells on the Nano patterned substrate was observed by the microscope (Nikon E600, Tokyo, Japan). FCPC areas on various size substrates at the single cell level are analyzed. The areas were quantified from the images in ImageJ. The formation of FCPCs aggregates was examined at 1, 5, 8 hours. The dynamics of aggregate formation after seeding till 20 hours was recorded by the live cell imaging. For live cell imaging, the temperature was set to 37°C by a microscope heating chamber and every 10 mins visualize the degree of self-assembly. N = 3 regions were imaged per condition.

### Scanning electron microscope (SEM)

In order to substrate and cell morphology, SEM was performed. Cells were fixed in 4% paraformaldehyde for a minimum of 10 min, followed by washing in PBS. Cell were incubated in a closed container of 100mM tetramethylorthosilicate (TMOS) solution in 1mM HCl at 40°C for 16–18 h, and allowed to air dry overnight. Cells were observed by SEM (JSM-6400Fs; JEOL, Tokyo, Japan) operated at a voltage of 5 kV.

### Immunocytochemistry

FCPCs were seeded onto Nano-particle coated coverslip for 5 h per condition. Cells were fixed with 4% paraformaldehyde. After blocking with 2% BSA in PBS, they were incubated with an anti-Tetramethylrhodamine antibody (1:1000; Sigma, ST. Louis, MO, USA) and anti-Ncadherin antibody (1:10; Abcam, Cambridge, MA, USA), anti-Integrin antibody (1:50; Abcam, Cambridge, MA, USA) for 1 h at room temperature.

### Centrifuge adhesion analysis

FCPCs were seeded in each condition 35mmΦ dish to measure the adhesion force. 1 h and 5 h after seeding, Unattached cells are washed with 1x PBS and counted. Add the 1x PBS (Dulbecco’s phosphate-Buffered Saline, Invitrogen) in a 35mmΦ dish and centrifuge at 3000rpm for 3min. The desorbed cells were washed; the cells still attached to the dish were described for 5 min using 0.05% trypsin-EDTA (Gibco BRL). Then, the number of adhesion cells in counted to calculate the adhesion percentage of the cells.

### Aggregation assay

FCPCs were seeded in each condition well with chondrogenic medium. To test the effects of cell adhesion, the following was re-suspended in chondrogenic medium: 1mM EGTA (Sigma). At 5 h after seeding, the effects of aggregate on the chondrogenic medium (+EGTA) observed by the microscope.

### Histological evaluation

For chondrogenic differentiation, 5×10^5^ cells were centrifuged at 500g 10 min, and the cell pellet was cultured. After 7 d of induction with chondrogenic medium, the samples were fixed with 4% formaldehyde (Duksan Chemical) and embedded in paraffin wax. Sections with a thickness of 4 μm were prepared and stained with Safranin O, hematoxylin and eosin (Sigma).

### Volume measurement of pellet culture

Total volume of aggregates construct was measured by a model 1076 X-ray micro-CT (SkyScan, Kontich, Belgium). Scanning was performed at a resolution of 15 μm pixel size (I = 190 μA, E = 40 kV, Filter = No filter). Scanned images were reconstructed with the software program (NRecon v.1.6.3.0, DataViewer v.1.4.4.). At 2, 4, and 7 d, aggregates were compared with the detected volume.

### Statistical analyses

The quantitative data were expressed as the mean ± standard deviation (SD) from at least three independent experiments. Statistical significance was analyzed by a one-way analysis of variance (ANOVA) followed by a Tukey–Kramer post-hoc test. A value of P < 0.05 was considered to be statistically significant (*P < 0.05, **P < 0.01 and ***P < 0.001).

## Results

### The morphology of single FCPC on silica nano-patterned substrate

The overall homogeneity of substrate coated with Nano-particles of different size was proved using scanning electron microscope (SEM) (Fig. 1). FCPC areas on various substrates at the single cell level are analyzed (Fig. 2A, B). Single FCPC is shown cell area larger, according to increasing particle size.

**Fig. 1.**
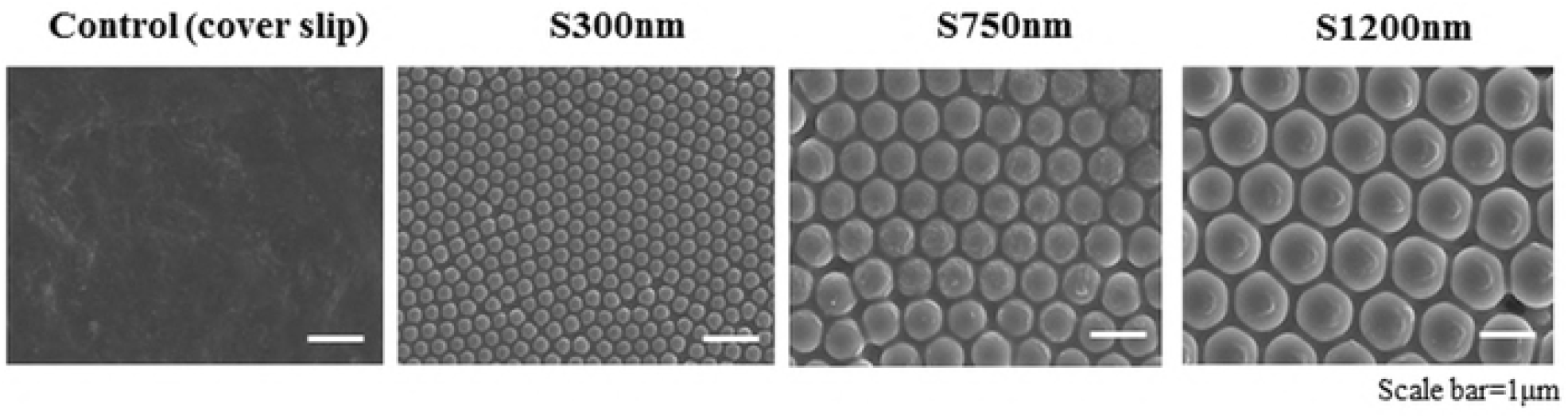
Scanning electron microscope (SEM) image of large surface areas displaying silica nano-patterned substrate. Image insets show the detail structure of silica 300nm (S300nm) and silica 750nm (S750nm), silica 1200nm (S1200nm).

**Fig. 2.**
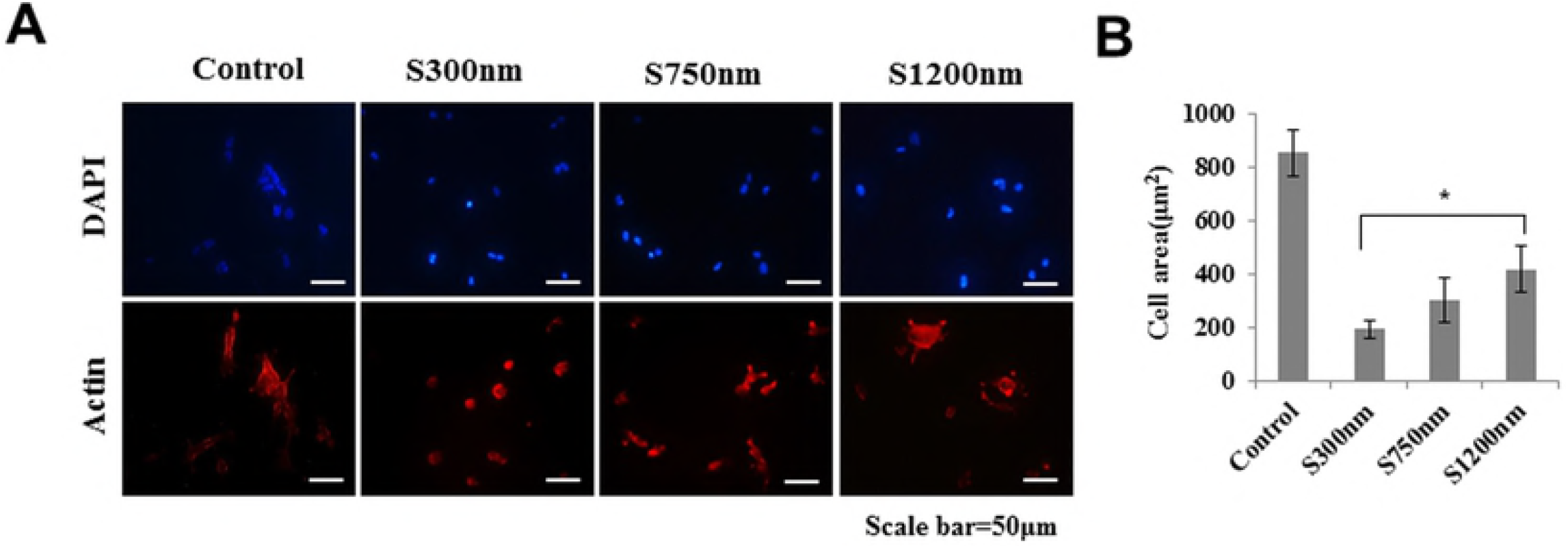
The morphology of single fetal cartilage-derived progenitor cells (FCPCs) by silica nano-particle size. (A) Single cell was fluorescently stained. (B) Single cell spreading area on various particle size surfaces.

### Aggregation formation of FCPCs by substrate coated Nano-particle size

By using a microscope, we visualized FCPCs in various substrates are shown (Fig. 3A). 5 h after seeding, FCPCs morphology is spread significantly increased, the larger the Nano-particles size (Fig. 3A, B, Supplementary Fig. 1A, B). When 10 h after seeding, Cells on the S300nm substrate is formed multi-spheroids and on the S750nm is formed mass-aggregation, in case of S1200nm is formed widely spreading like as a conventional cell culture plate. The manufactured 3D aggregates construct (S1200nm could not be collected a cell, and could not manufacture it) were examined up to 7 d by measuring their volume using μ-CT (Fig. 3B). Change in volume of construct at 1, 4, and 7 d were 5.18, 6.62, 0.86 mm^3^ (S300nm, S750nm, pellet) at 1 d and 5.24, 6.68, 1.33 mm^3^ at 4 d and 6.88, 8.12, 1.68 mm^3^ at 7 d. Aggregates at S750nm condition maintain large size for 7 d.

**Fig. 3.**
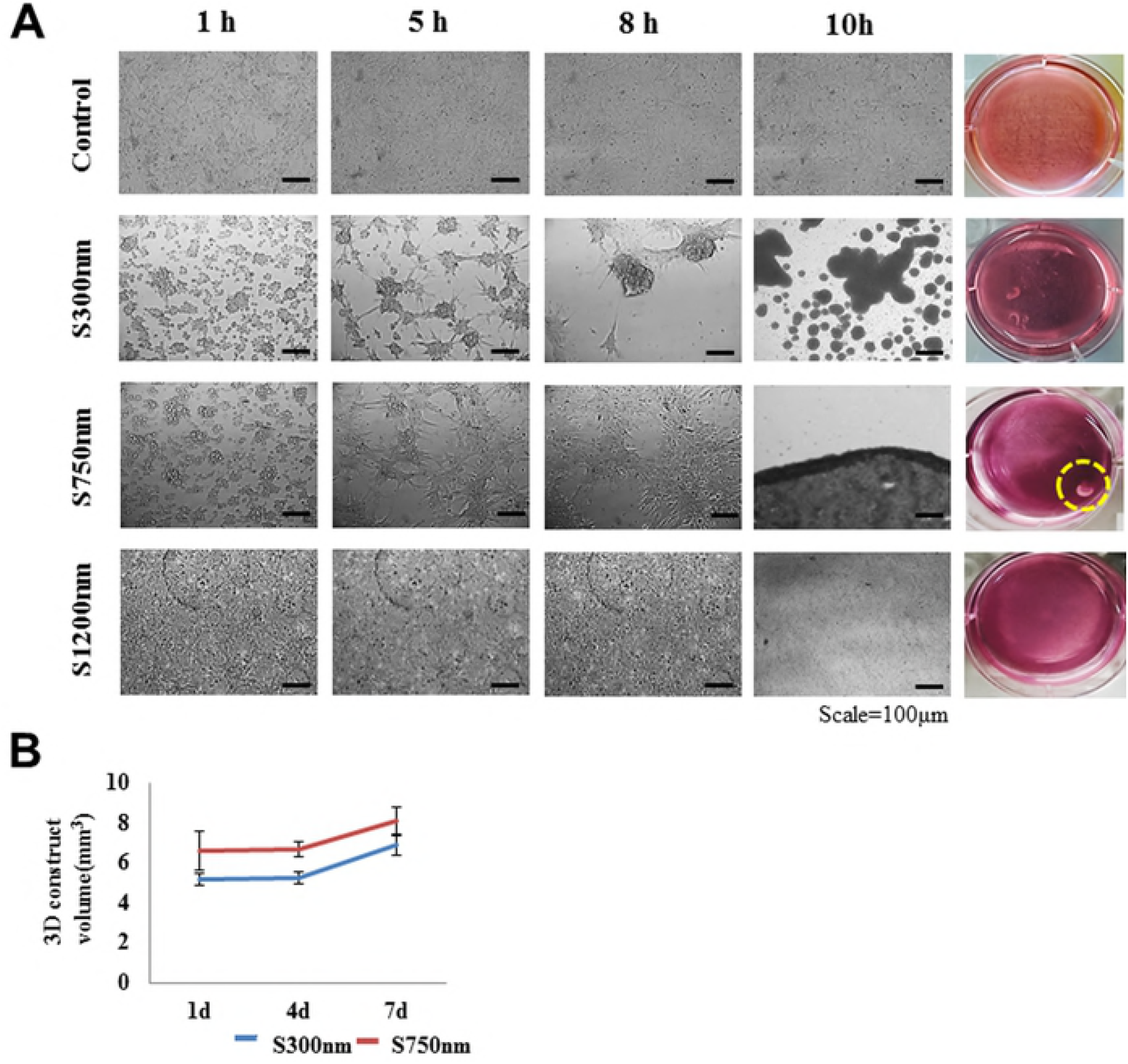
3D articular cartilage construct is formed by Silica nano-patterned substrates. The morphologies of FCPCs on Nano-patterned plate with different silica size at each time (1, 5, 8, and 10 h). (B) The volume of each construct was measured using a custom-made device and software as described previously. Data were presented as mean values with standard deviation (SD) from four independent experiments (*n*=5).

### The Chondrogenic potential of 3D aggregates constructs of FCPCs using Nano-patterned substrate

Cartilage regeneration ability to construct was evaluated at 1, 7 d by Safranin-O staining, hematoxylin and eosin (Fig. 4A). At 7 d, GAG accumulation of the construct was strongly by safranin-O in construct from S750nm. At 7 d, lacuna formation by hematoxylin and eosin was similar in both S300nm and S750nm conditions (Fig. 4B).

**Fig. 4.**
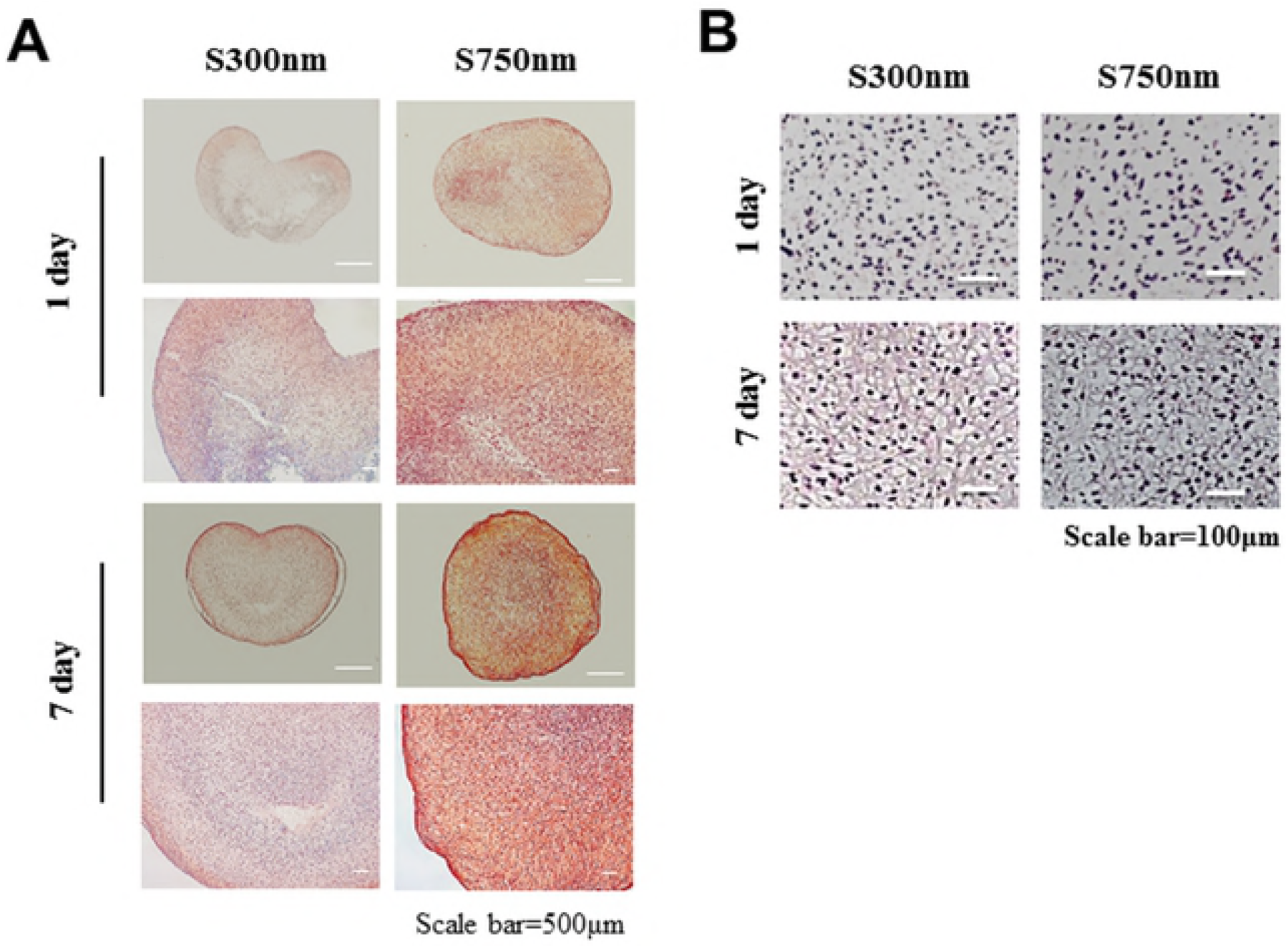
Histological of (A) S-O staining, (B) hematoxylin and eosin staining of cartilage from each gruoop at 1, 7 days.

### The role of cell adhesion molecules in FCPCs aggregation on various substrates

The expression differences of integrin, which are cell-material adhesion molecules, were analyzed by immunocytochemistry (Fig. 5A). The expression of Integrin β1 is the largest at S1200nm substrate. It is the results that cell-material interaction is more involved at S1200nm. Also, the centrifuge adhesion technique was used to determine the relative adhesion force on various substrates in 1 h and 5 h after seeding. A bar graph shows percentage of single FCPC adhered to the surface of the 35mmΦ dish when centrifuge conditions at 3000rpm and 3 min (Fig. 5B).

**Fig. 5.**
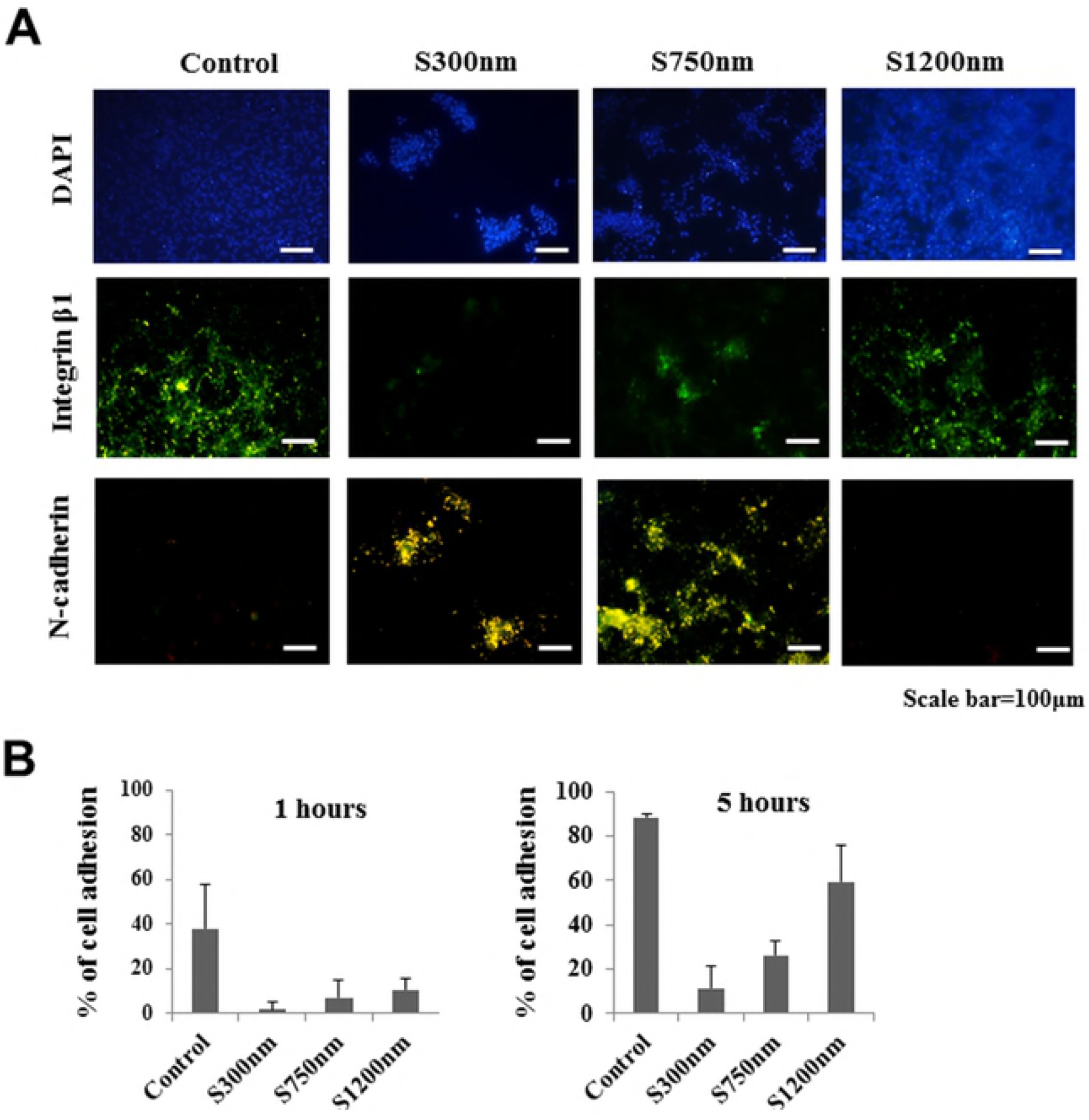
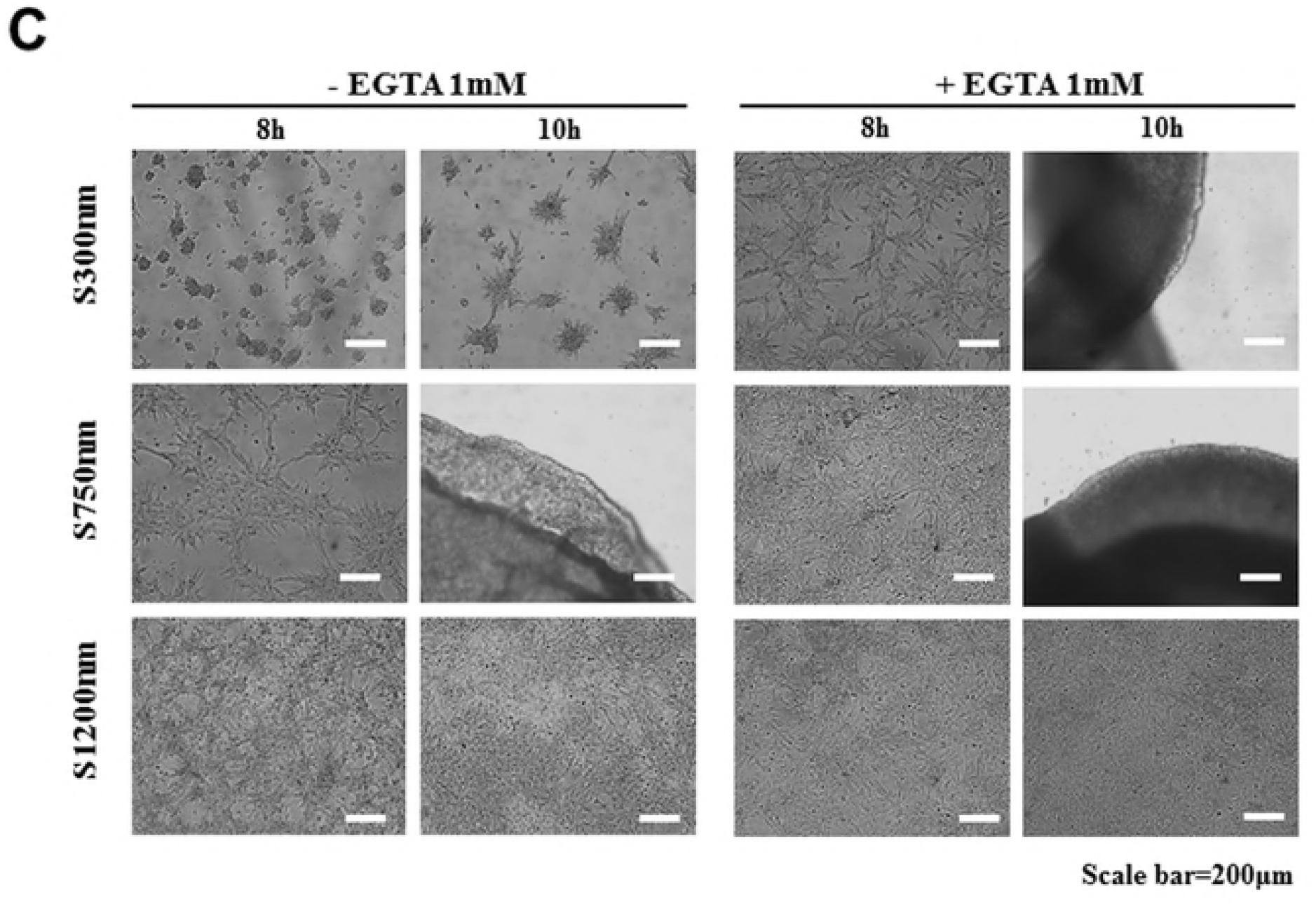
Differences in cell interaction in FCPCs aggregation by Nano-patterned substrate. (A) FCPCs cultured on various particle size surfaces for 5 h and stained with monoclonal anti-Integrin β1 (green, cell-matrix adhesion), N-cadherin (green, cell-cell adhesion). Scale bar = 100 μm. (B) Percentage of adhesion cell number after assay between 1 h and 5 h. The adhesion assay used rotating spinning disc at 3000rpm for 3min,*p<0.05, **p<0.01 and ***p<0.001. (C) Cell aggregation assay was performed in the presence of EGTA reagents. The morphologies of FCPCs cultured on various particle size surface in the presence of 1mM EGTA (Ca^2+^ chelating agent) in the chondrogenic medium.

The expression differences of N-cadherin, which are cell-cell adhesion molecules, were analyzed by immunocytochemistry (Fig. 5A). The expression of N-cadherin appears to be the largest at S300nm substrate. It is the results that cell-cell interaction is more involved at S300nm. Also, the FCPCs suspensions were treated with EGTA that chelate Ca^2^+ (Fig. 5C). EGTA chelate calcium that is an inhibitor for calcium-dependent, cadherin-mediated cell-cell interaction. FCPCs treated with EGTA, widely spread without forming spheroids (S300nm) or the aggregation time was delayed (S750nm). On the other hand, there was no significant difference in S1200nm.

## Discussion

In the current study silica bead was coated into the well of a 6-well tissue culture plate (Fig. 1). Silica beads are chemically inert and extremely stable; they provide a greater surface area for the adhesion of cells and biomolecules (Fig. 2) [24]. In this study, we manufactured three-dimensional cartilage construct utilizing FCPCs cultured on silica nano-patterned substrate through particle diameters of 750 nm (Fig. 3). The process of aggregation formation for cells on nano-patterned substrate appears to be different from each study. Embryonic stem cells (ESC) were not spreading on the silica colloidal crystal 400nm substrates to the same extent as they did on glass cover slips, the reduction of cell-substrate contacts promotes ESC differentiation [25]. Bovine endothelial cells tend to spread less with larger particle size and fewer focal adhesion complexes. In contrast, Mouse preosteoblasts showed increasing their area of contact with larger particle size and many focal adhesion complexes [15]. These results suggest that there exists a critical dimension of Nanoparticles size for affecting cell behavior on shape, differentiation, adhesion and these effects can know that cell type specific. Actually, Different results were confirmed from the silica nanopatterned substrate used in this study: MSCs tend to spheroid with larger particle size (350-1200 nm), but it is the multi-spheroids, not one large aggregation (Supplementary Fig. 2). The FCPCs is forming larger aggregations with 300 nm, 750 nm particle size, but not at 1200 nm particles size. According to the result of this study, 750 nm particle size, which forms one large self-aggregation, was most useful, depending on the purpose of this study.

Culturing cells in 3D narutally enhance cell growth, differentiation and function. Although these 3D culture methods are used in many studies, they are limited in that they are not large size enough to be used in clinical applications. Traditional methods such as the use of low attachment plate and hanging drop, pellet culture methods, etc., show that the final aggregation size is very small in micro units and there is a limit on the number of cells for constructing the compacted cell structure [26],[27],[28],[29]. In terms of methods pellet culture and rotary culture and orbital shaker have the limitation that external force is needed in the process[10],[29]. Also, the aggregates formation by enzymatically detached cells is likely to break the natural ECM[30]. In our results, the spontaneous aggregation phenomena seen did not require any external force or enzyme digestion for cell detachment, and formed one large mass-aggregation keeping their naturally networks intact without breaking ECM (Fig. 4). Thus, this aggregation method through nano-patterned substrate suggested herein might lead to a great improvement of general methods. In particular, it is a very superior advantage for use in cartilage tissue engineering.

Spheroid self-assembly consisted of a series of events involving that cell-matrix forces through focal adhesion contacts are generally present in spreading cells. At this stage, the cell forms a monolayer like cell sheets. These cell monolayers undergo contraction through cell-cell contacts force [21]. A number of studies have confirmed cell adhesion molecule to analyze the cell aggregation mechanism. Among them, we have chosen to N-cadherin involved in cell-cell adhesion and Integrin involved cell-materials adhesion (Fig. 5A). Cadherin and Integrin are two of the best-studied classes of adhesion receptors. The interaction between integrin and cadherin can direct the localization of forces within cell aggregates through the tensional changes in the actin cytoskeleton[31]. In this studys, the 300 nm substrate that formed multi-spheroids and the 1200 nm substrate that showed spreading were due to the cell-cell adhesion force (via N-cadherin) and cell-substrate (via Integrin) force, the 750 nm substrate that formed the mass-aggregation can be interpreted as the result of cell monolayer formation through cell-substrate force followed by cell-cell contact force contraction.

In this study, the spatial distribution of the nanoscale is an important feature that has a significant effect on cellular material interactions [32]. A thorough review of the effects of feature spacing on integrin adhesion suggests that featured can interfere with integrin binding in any size rage, which is consistent with decreasing cell spread [33]. The aggregation of FCPCs on Nano-patterned substrate in the presence of EGTA is shown Fig. 5C. N-cadherin mediated homotypic cell-cell adhesion in a ca^2+^ dependent manner. To tease out the potential roles of N-cadherin, the treated with EGTA that chelate Ca^2^+ [34],[35],[36]. In the results, the possible correlation between aggregation time delay and inhibition of N-cadherin by Ca^2+^ reduction was diminished cell contraction due to inhibition of cell-cell contacts. But, the 1200 nm substrate showed no significant differences between presence and absence of EGTA. It is conceivable that this would be the condition which N-cadherin is less affected. Based on the results, cell adhesion force is affected by the regulation of substrate coated Nano-particle sizes.

## Conclusions

Collectively, our data can judge that FCPCs aggregation using Nano-patterned substrate of tissue engineered constructs has a beneficial effect on 3D culture for cartilage regeneration compared to conventional 3d culture methods. Self-aggregation phenomena in our experiments can focus on events corresponding to the possibility of Self-aggregation called condensation of mesenchymal cells for chondrocyte differentiation. However, the constructs produced in this study were confirmed only the histological evaluation whether it’s effective in cartilage regeneration. Therefore, it is necessary to verify that this construct actually has a cartilage tissue regeneration potential afterwards.

## Supplementary Figure

**S Fig. 1.**
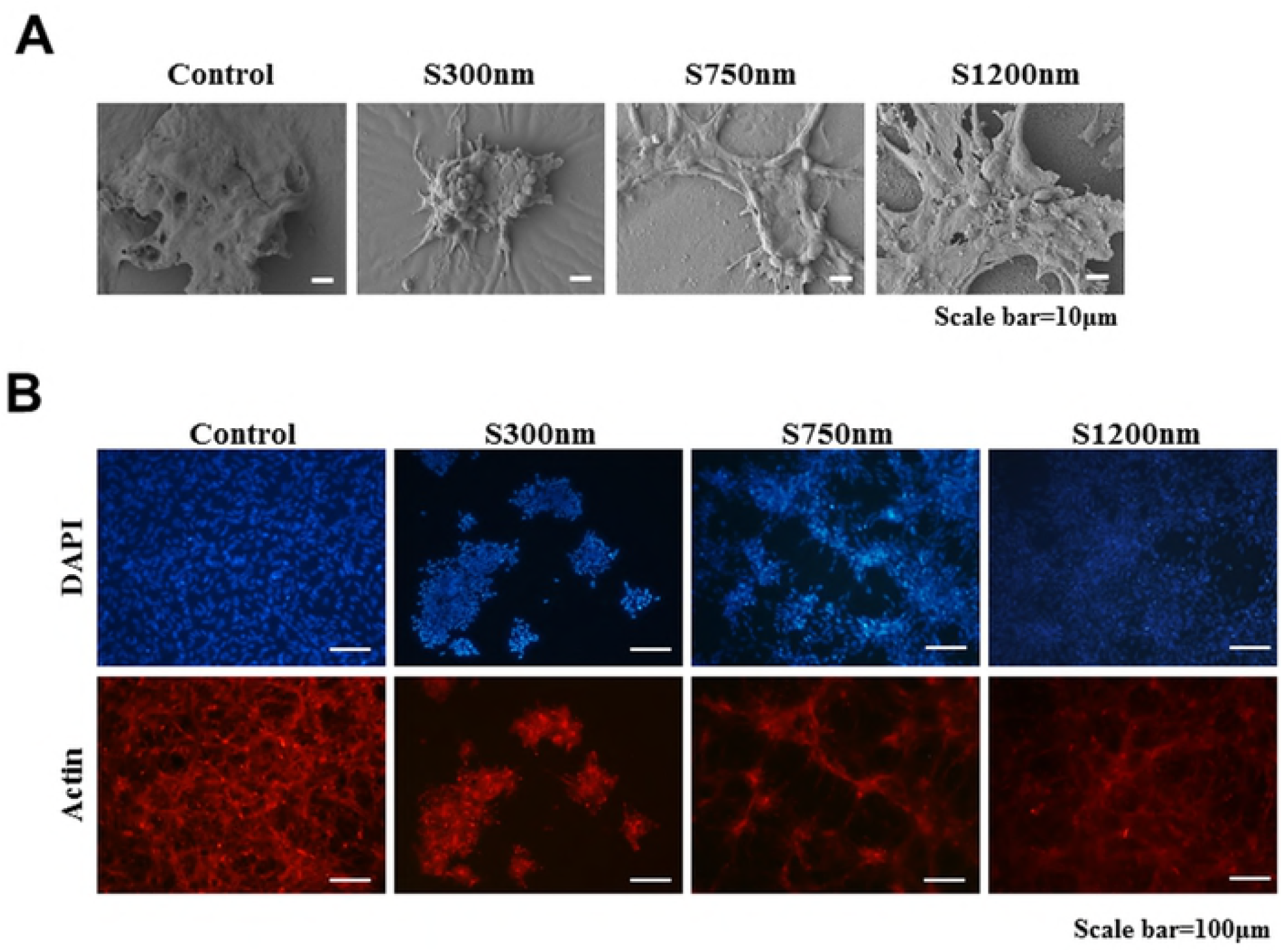
FCPCs aggregation morphology by particle size. 5 h after seeding, (A) Scanning electron micrographs (SEM) of cells cluster on S300nm, S750nm, S1200nm. (B) F-actin (red) and nucleus (blue) of cells cluster was fluorescently stained.

**S Fig. 2.**
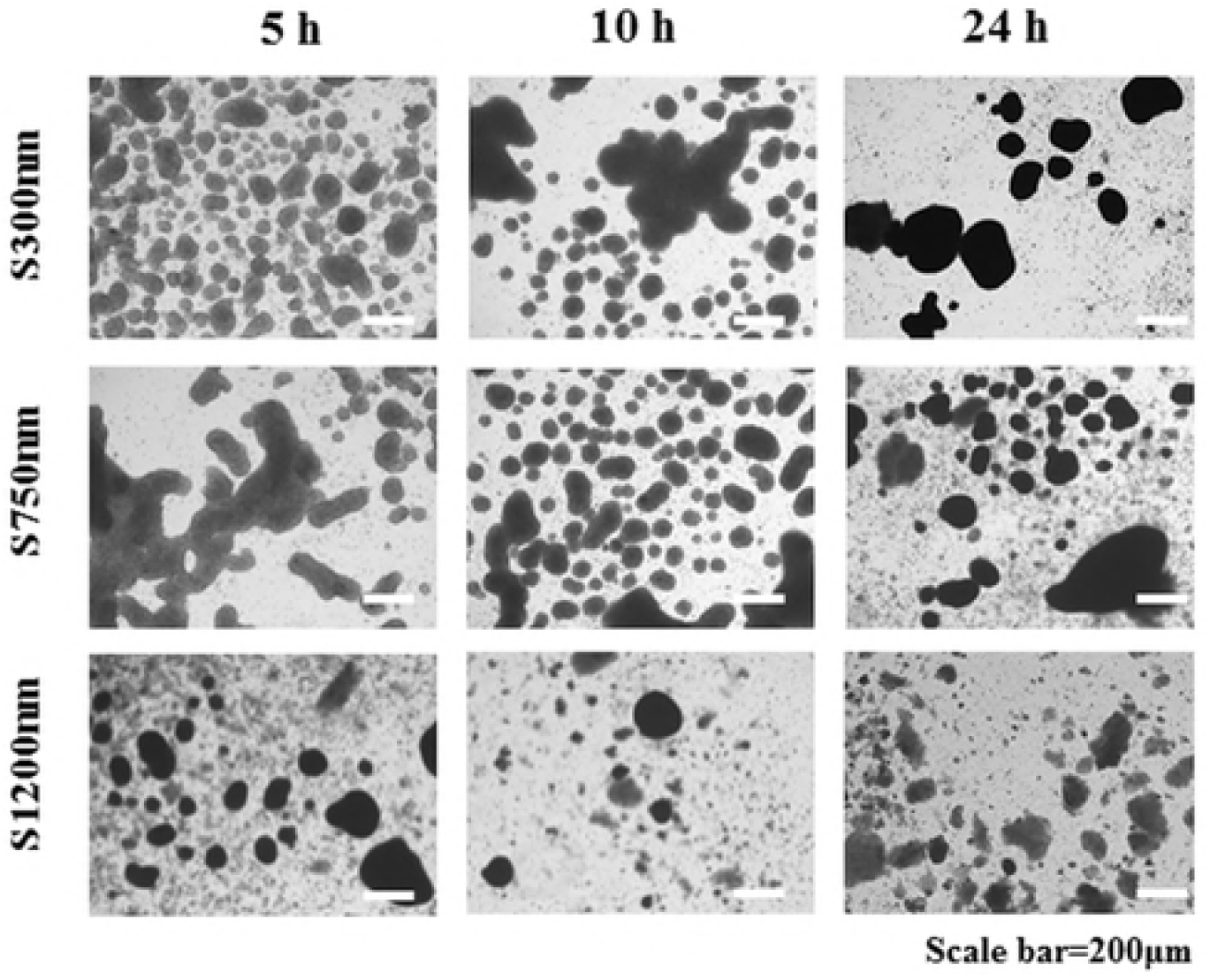
The morphologies of mesenchymal stem cells (MSCs) on nano-patterned substrates with different silica size at each time (1, 10, and 24 h).

